# A new subspecies of gray wolf, recently extinct, from Sicily, Italy (Carnivora, Canidae)

**DOI:** 10.1101/320655

**Authors:** Francesco Maria Angelici, Lorenzo Rossi

## Abstract

A new endemic subspecies of gray wolf from the island of Sicily (Italy) is described. While usually considered extinct before 1940, there’s some evidence it may have survived up to 1970. This wolf was widespread throughout the island and characterized by a smaller size and a paler coloration than the Apennine wolf *(Canis lupus italicus)* from Central-Southern Italy.

This subspecies is described from a mounted specimen (the holotype) including also a separate skull stored at the Museo di Storia Naturale ‘La Specola’, Università di Firenze, Italy. The three paratypes are: a) a mounted specimen stored at the ‘Museo Regionale Interdisciplinare di Terrasini’ in Terrasini (PA), Italy, b) a mounted specimen stored at the Museo di Zoologia ‘Pietro Doderlein’, Università di Palermo, Palermo, Italy, c) a mounted specimen stored at the ‘Museo Civico Baldassarre Romano’ in Termini Imerese (PA), Italy.

This new subspecies is described as *Canis lupus cristaldii* **subsp. nov**. We suggest ‘Sicilian wolf’ as common name for this new *taxon*.

## Introduction

The gray wolf *(Canis lupus)* is a widely distributed species native to many Eurasian and American regions (Mech and Boitani 2010). Currently there are at least forty-four subspecies described (see Wozencraft 2005) several of which are however disputed. Among these is the Apennine wolf *Canis lupus italicus* Altobello, 1921, widespread in the Italian peninsula and morphologically and genetically distinct from all other European populations (Nowak and Federoff 2002; Montana *et al*. 2017).

In the past *Canis lupus* was also present in Sicily, the largest island of the Mediterranean Sea, located south of the Italian Peninsula (about 37°45’0 N; 14°15’0 E). It was especially common in the mountains around Palermo, the woods around Mt Etna, the Peloritani, Nebrodi, Madonie and Sicani Mountains and Ficuzza Wood; the species was also present further south on the Erei and Iblei Mountains, where it has been recorded until 1928 (La Mantia and Cannella 2008).

The Sicilian wolf is usually considered extinct in the early decades of the twentieth century, but there is no unanimity on the exact date. The last confirmed specimens were shot near Bellolampo (PA) in 1924, but there are several reports of wolves killed between 1935 and 1938 near Palermo. Furthermore, there are several sightings of wolves in Sicily between 1960 and 1970 and some of them seem convincing (Angelici *et al*. 2016a).

The main cause of extinction of the Sicilian wolf appears to have been human persecution, due to alleged damage to livestock (eg. Minà Palumbo 1858a; 1858b; Chicoli 1870). This may have been due the extinction of the wild ungulates on which the Sicilian wolf fed due to an environmental crisis that starting on the island at the end of the Norman Period (ca 1198 CE) (Bresc 1983).

To date, remains of Sicilian wolves are extremely scarce: no more than 7 specimens (represented by skins, stuffed individuals, skulls, etc.) are known in Italy, in the ‘Museo Regionale Interdisciplinare di Terrasini’, Terrasini (PA), in the Museo di Zoologia ‘P. Doderlein’, Università di Palermo, in the Museo di Storia Naturale ‘La Specola’, Università di Firenze, and in Museo Civico ‘Baldassarre Romano’, Termini Imerese (PA). Moreover, there exists only a single picture of a live Sicilian wolf (Fig. 1).

**Fig. 1.**
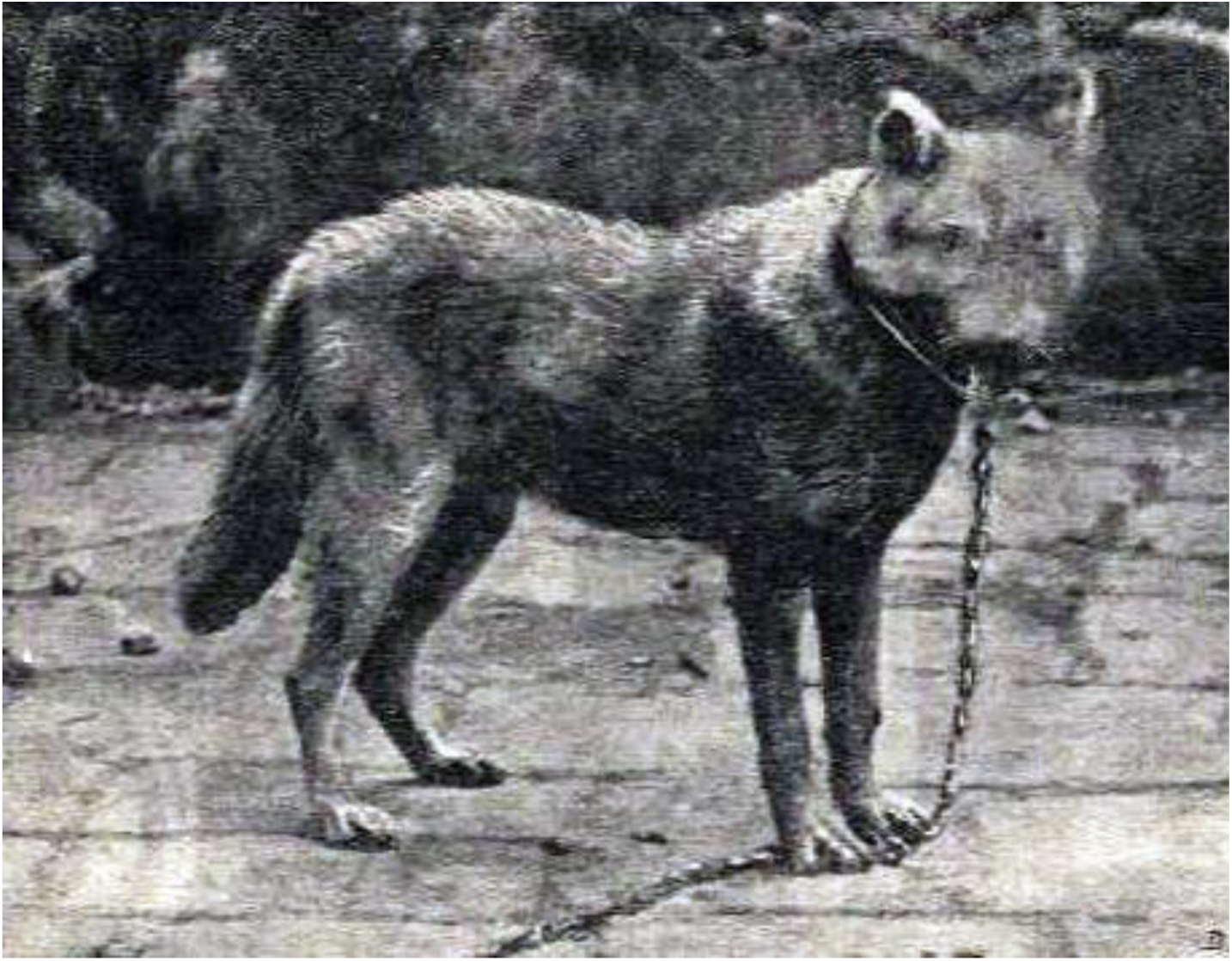
Sicilian wolf held in captivity in the late XIX century. This image turns out to be the only image of a live Sicilian wolf (from Migneco, 1897).

The complete specimens available had some peculiarities compared to the Apennine wolf. Among these the smaller size and the paler coat colour, which have already been noticed by some authors of the past (eg. Minà Palumbo 1868; Migneco 1897).

## Materials and methods

A total of four adult specimens were studied: the holotype, a mounted male specimen plus its separate complete skull, and three paratypes, three mounted specimens (all males). Although Sicilian wolf disappeared only recently there are very few specimens available and some of them are in poor conditions. We were also able to find three other specimens: another skin and two juveniles. However we chose not to use these in our description.

Repositories of described or cited specimens are: Museo di Storia Naturale ‘La Specola’, Università di Firenze, Florence, Italy; ‘Museo Regionale Interdisciplinare di Terrasini’, Terrasini (PA), Italy; Museo di Zoologia ‘Pietro Doderlein’, Università di Palermo, Palermo, Italy; Museo Civico ‘Baldassarre Romano’, Termini Imerese (PA), Italy.

Body measurements (n= 4) were taken using a tape measure to allow flexibility on mounted specimens, with an accuracy of 0.1 cm

A total of 10 measurements were taken on the skull of the holotype: 7 measures from the skull, and 3 from the mandible.

The skull measurements were taken using a Vernier gauge with accuracy of 0.1 mm

The skull belongs to an elderly individual and several teeth are missing, while those still present are visibly worn and blunt. Only measurements that can be obtained with the highest precision allowed by our instruments were thus taken.

We compared skull and body measurements of the Sicilian wolf with the respective measurements of *C. l. italicus* using data available in literature (Siracusa and Lo Valvo 2004; Berté 2013; Altobello 1921; Ciucci and Boitani 2003). We have excluded from the comparative skulls the specimens with deciduous teeth or teeth not yet completely erupted (cf. Siracusa and Lo Valvo 2004; Berté 2013). We used only average measurements for sexually mature, adult individuals. (Altobello 1921; Ciucci and Boitani 2003).

In order to sort the multidimensional data, we developed a Principal Component Analysis (PCA) of body measurements from a sample of twenty Apennine wolves (9 MM, 11 FF) and a sample of four males Sicilian wolves using the software ‘R’. Then a cluster based on Euclidean distances between the points of the PCA to reconstruct the phenetics relationship was made using the Ward Method (1963). This method was chosen because it recognizes groups that minimize intra-group variance (more morphologically homogeneous).

## Systematic taxonomy

**Carnivora Bowditch, 1821**

**Caniformia Kretzoi, 1938**

**Canidae Fischer, 1817**

***Canis* Linnaeus, 1758**

***Canis lupus* Linnaeus, 1758**

***Canis lupus cristaldii* subsp. nov. (Sicilian wolf)**

**Holotype**. Mounted specimen of an old adult male, labeled M1891-Coll.652-1884 (Fig. 2). Skull C11875 (Fig. 3). The wolf was killed on 17^th^ July 1883, at Vicari (PA). Collected by Mr Costantino Ciotti. Museo di Storia Naturale, Sezione di Zoologia, ‘La Specola’, Università di Firenze, Florence, Italy

**Fig. 2.**
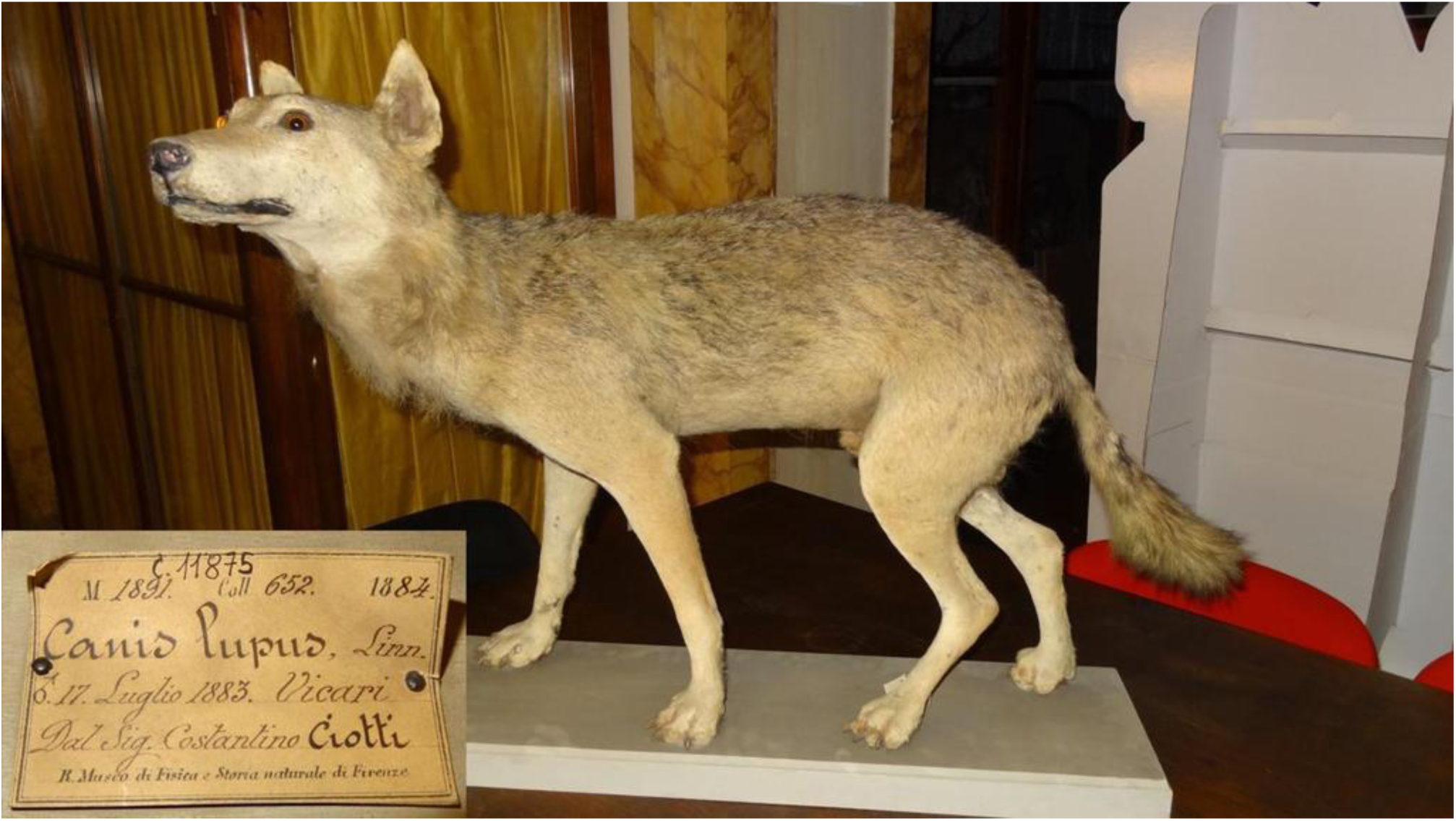
*Canis lupus cristaldii* subsp. nov. Holotype. Mounted specimen of an old adult male, labeled M1891-Coll.652-1884. Museo di Storia Naturale, Sezione di Zoologia ‘La Specola’, University of Florence, Florence, Italy.

**Fig. 3.**
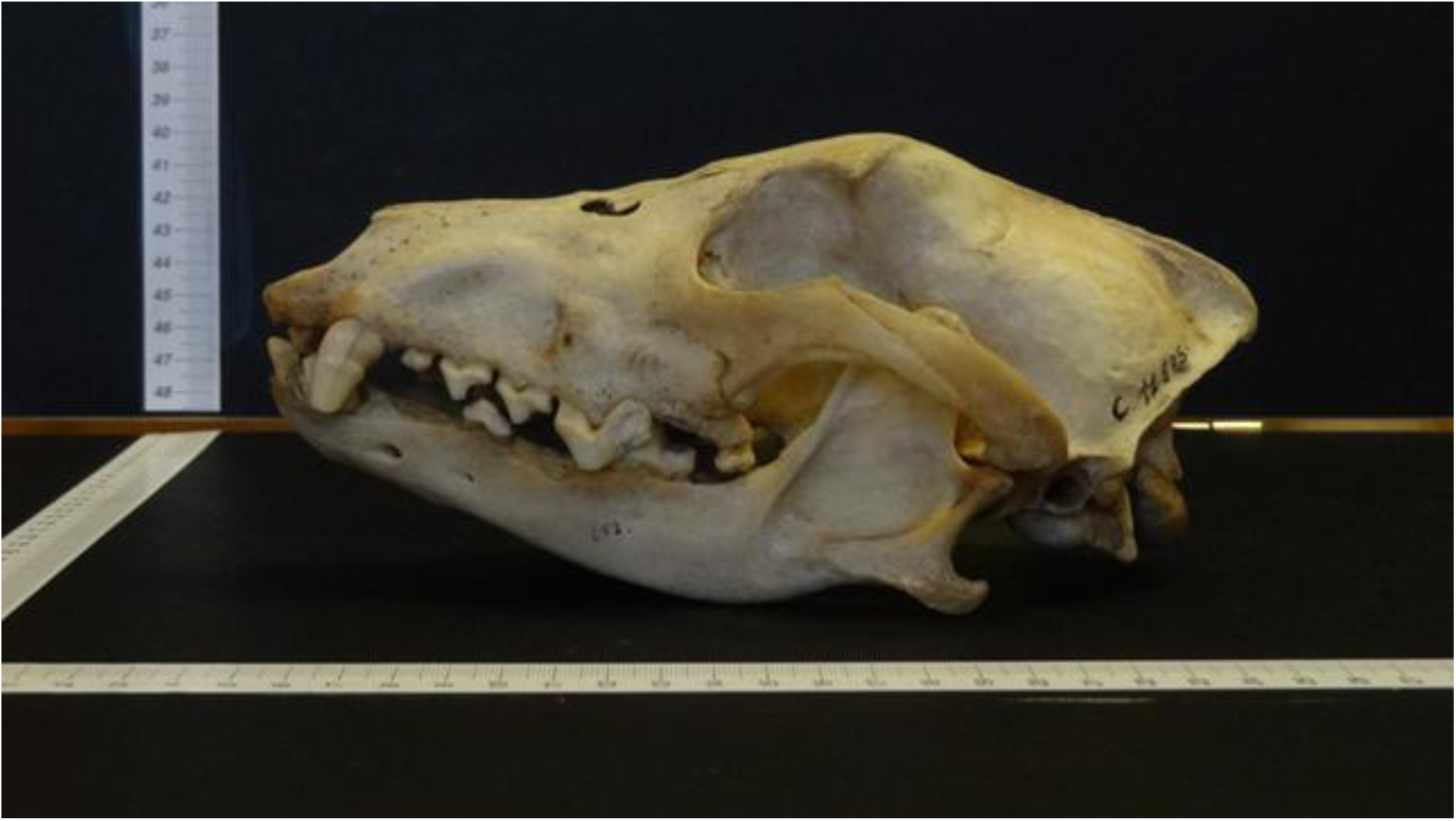
*Canis lupus cristaldii* subsp. nov. Holotype. Skull number C11875. Museo di Storia Naturale, Sezione di Zoologia, ‘La Specola’, Università di Firenze, Florence, Italy.

**Paratypes**. N. 1 Mounted specimen of an adult male, inventory number 9263 (Fig. 4). This individual was caught in Sicily (unknown location), date unknown. Donated by ‘Centro Studi e Ricerche’, C.S.I.’, Trapani, Italy. Museo Regionale Interdisciplinare di Terrasini (PA), Italy

**Fig. 4.**
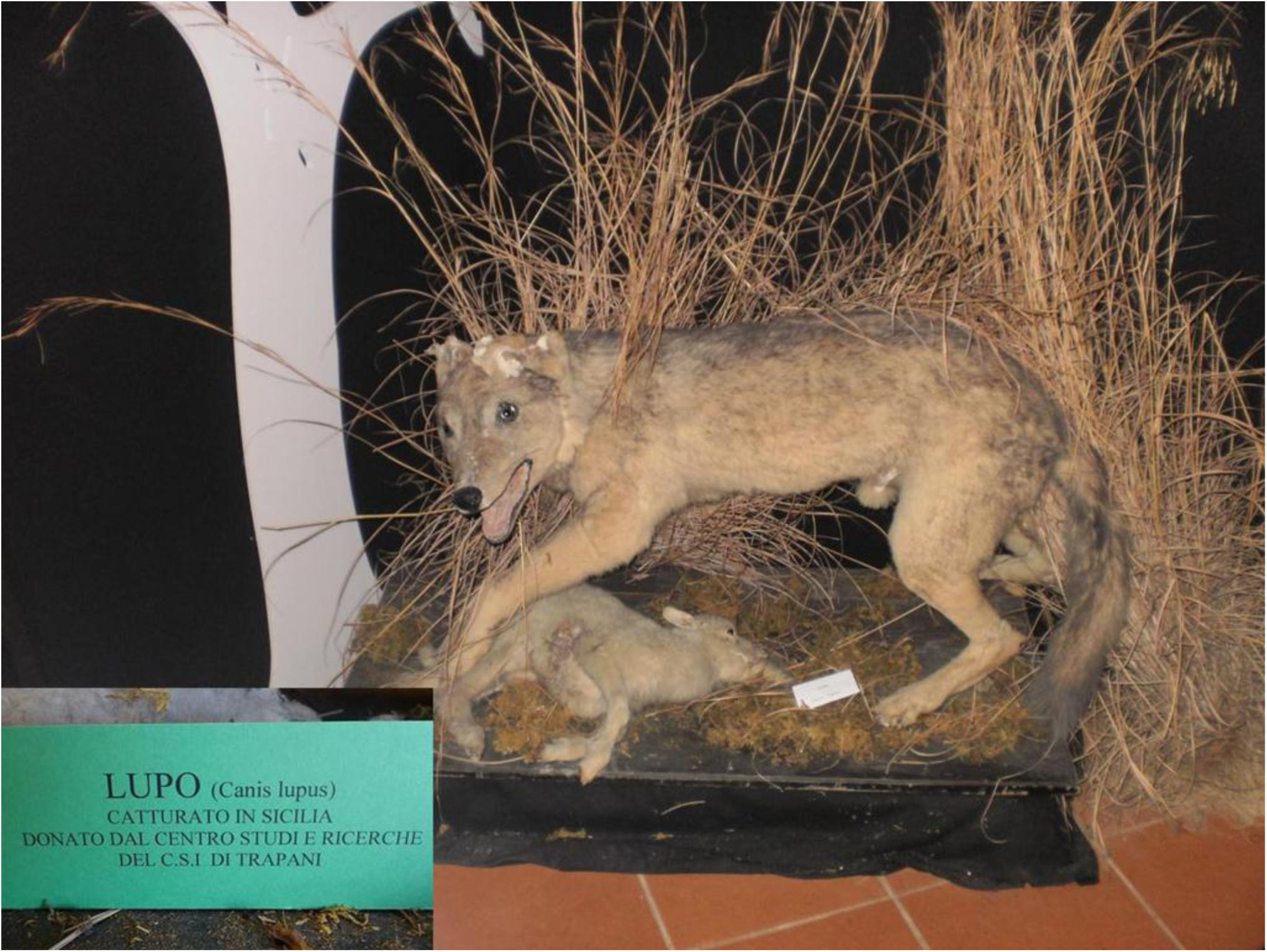
*Canis lupus cristaldii* subsp. nov. Paratype n.1. Mounted specimen of an adult male, inventory number 9263. Museo Regionale Interdisciplinare di Terrasini (PA), Italy.

N. 2 Mounted specimen of an adult male, labeled M/18 (Fig. 5). This individual was killed in Sicily (unknown location), date unknown. Museum of Zoology ‘Pietro Doderlein’, Università di Palermo, Palermo, Palermo, Italy

**Fig. 5.**
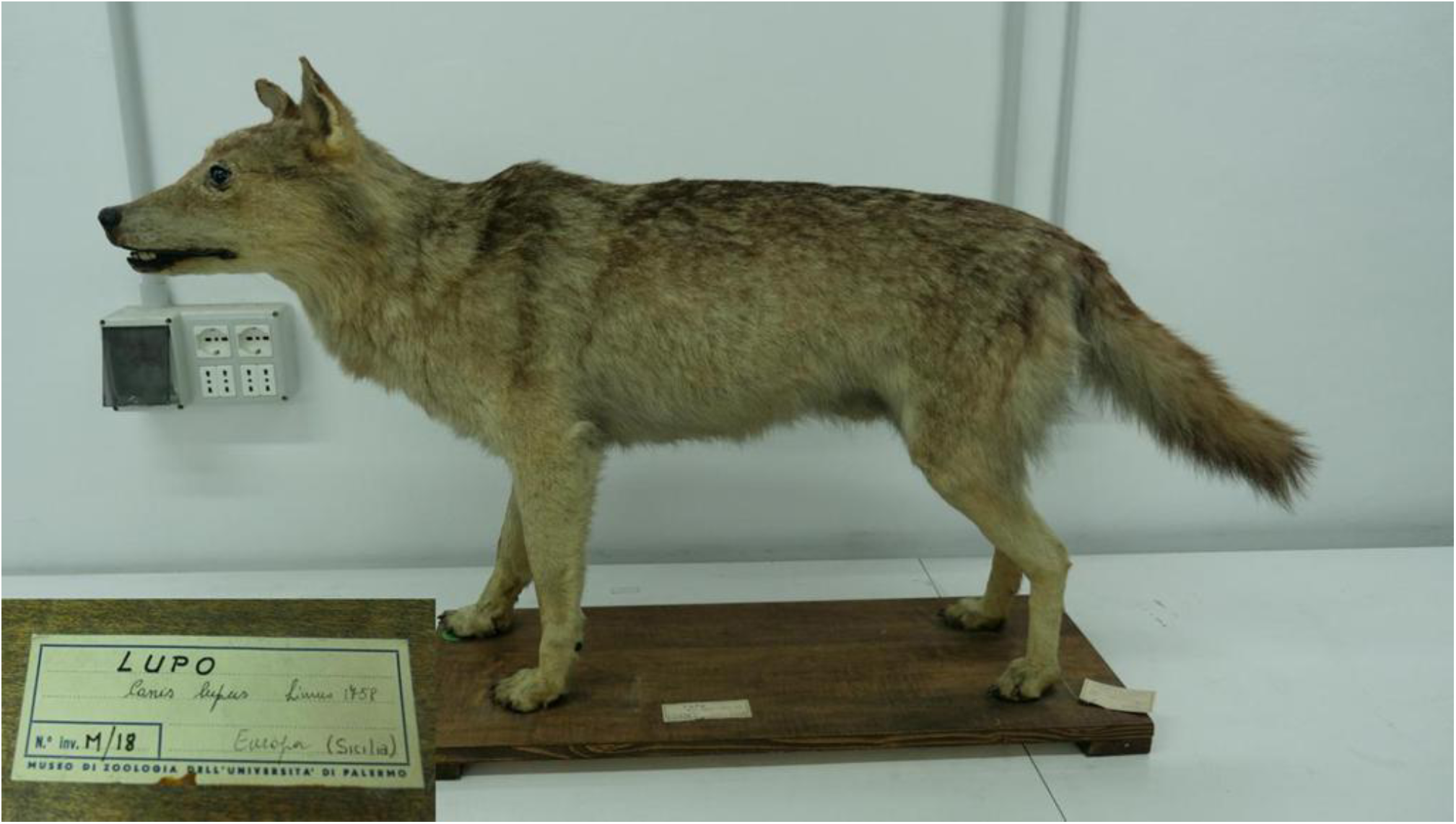
*Canis lupus cristaldii* subsp. nov. Paratype n.2. Mounted specimen of an adult male, labeled M/18. Museum of Zoology ‘Pietro Doderlein’, University of Palermo, Palermo, Italy.

N. 3 Mounted specimen of an adult male, labeled 3 (Fig. 6). This individual was most likely shot on Monte San Calogero near Termini Imerese (PA), date unknown but most likely in the last years of nineteenth century. The specimen was acquired by the Museum in 1969. Museo Civico ‘Baldassarre Romano’, Termini Imerese (PA), Italy

**Fig. 6.**
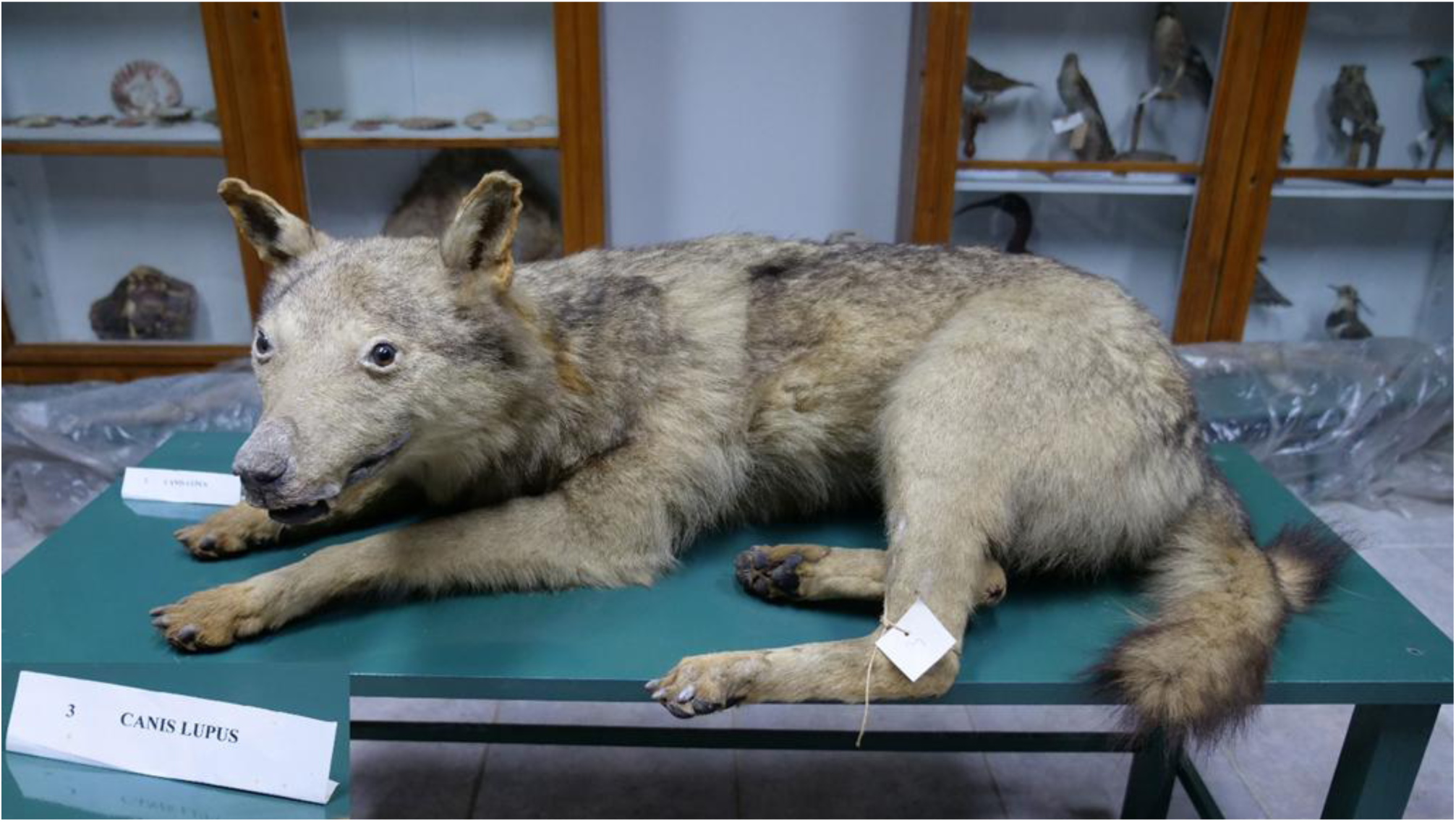
*Canis lupus cristaldii* subsp. nov. Paratype n.3. Mounted specimen of an adult male, labeled 3. Museo Civico ‘Baldassarre Romano’, Termini Imerese (PA), Italy.

**Etymology**. This taxon is dedicated to Professor Mauro Cristaldi (1947-2016), an Italian mammalogist and professor of Comparative Anatomy, born in Rome from a family of Sicilian origins. He was very attached to Sicily and he devoted his life to the study of Italian mammals, mostly rodents. He died suddenly in the summer of 2016.

**Type locality**. Vicari (Palermo Province), 640 m a.s.l., near ‘Bosco della Ficuzza’, NW Sicily (37°51’0.36” N, 13°34’0.72” E)

**Distribution**. In the past the species was widespread all over the island, especially around Palermo, woods surrounding Mt Etna, Peloritani, Nebrodi, Madonie and Sicani Mountains, and the Ficuzza Wood. It was also present further south, on the Erei and Iblei Mountains.

**Diagnosis**. The body is slender and sturdy, but overall proportionate. The legs are short. The small size of the Sicilian wolf place it among the smallest subspecies of *Canis lupus*, together with the endangered Arab wolf *Canis lupus arabs* Pocock 1934, and the extinct Japanese wolf *Canis lupus hodophilax* Temminck 1839 (see Pocock 1935).

To our knowledge there are only two works in literature containing measurements of Sicilian wolves (Galvagni 1837; Minà Palumbo 1868) but use of these sources is problematic. Galvagni (1837) uses inches as units of length, which we have considered *pouces* or Paris inches (1 Paris inch = 2.70 cm) for conversion in International System. Moreover, Galvagni (1837) does not specify if the height measurement is to ‘shoulder height’ or ‘withers height’ and we believe he may have included the head.

Minà Palumbo is also not clear in indicating the height, and he also does not specify if the ‘body length’ includes the tail or not. Considering the sizes reported it is probable he included tail measurements as well.

The body measurements of four *C.l.cristaldii* subsp. nov. specimens studied are in Table II.

**Table I.** Historic measurements of the Sicilian wolf

|  | **Measurements by Galvagni (1837) (n=?)** | **Measurements (mean) by Minà Palumbo (1868) (n = 6)** |
| --- | --- | --- |
| Head-body length | 3 feet 3 inches<br>(105.3 cm) | 125 (cm) (including tail ?) |
| Tail length | 9.5 inches<br>(25.65 cm) | 21 (cm) |
| Height (including head ?) | “more than 2 feet”<br>(more than 64.8 cm) | 63 (cm) |

**Table II.** *Canis lupus cristaldii* subsp. nov. body measurements

|  | Measurements taken on available specimens (n = 4) |
| --- | --- |
| Head-body length (cm) | (108; 110; 101; 102.5) |
| Tail length (cm) | (30.5; 43.5; 32; 33.5) |
| Shoulder height (cm) | (52.5; 53; 56.5; 56.5) |
| Ear length (cm) | (9; 8; 9.5; 7) |

The overall colour of the coat varies slightly from individual to individual and is generally paler than those displayed by other European wolf populations, particularly *C. l. italicus*. We believe the mounted specimens we used for this study did not suffer a significant discoloration. First, in XIX century sources the colour of the Sicilian wolf is described as very similar to that of the studied specimens for both living and recently dead adult individuals (eg Minà Palumbo 1869). Furthermore at least three of the four studied individuals (including the holotype) have been stored away from direct sunlight for many decades.

The background colour is very pale tawny, in some way reminiscent of the lion’s coat (‘lionato’ according to Minà Palumbo 1868). A good match in commonly available literature is Lion tawny, Dictionary of Color #C19A6B.

Older individuals are ashen in colour, almost dirty white (see also Minà Palumbo 1868). Dorsally and on the sides the coat is overall darker, with predominance of light tawny, dark grey, and lighter grey tones. The colour of the chest and the belly is light and varies from dirty white to pale yellowish cream. The throat, cheeks and inner parts of the limbs are also paler than the dorsal parts. The ears are light beige-tawny with some darker grey dorsally hairs and very light, almost white, internally hairs.

The base of tail is narrow, then widens and the hair becomes thicker, ending progressively in a pointed shape. The colour of the tail is usually darker at the base, sometimes tending to black at the end.

The legs are rather pale, between light beige and grey. The dark line on the forearms, typical of Apennine wolf (Altobello 1921), is absent or very slightly hinted (see Fig. 7). However, there is a very slight difference between the external and the internal colour of the legs: the latter is lighter. Intermediate hairs of the toes tend to ochraceous-tawny.

**Fig. 7.**
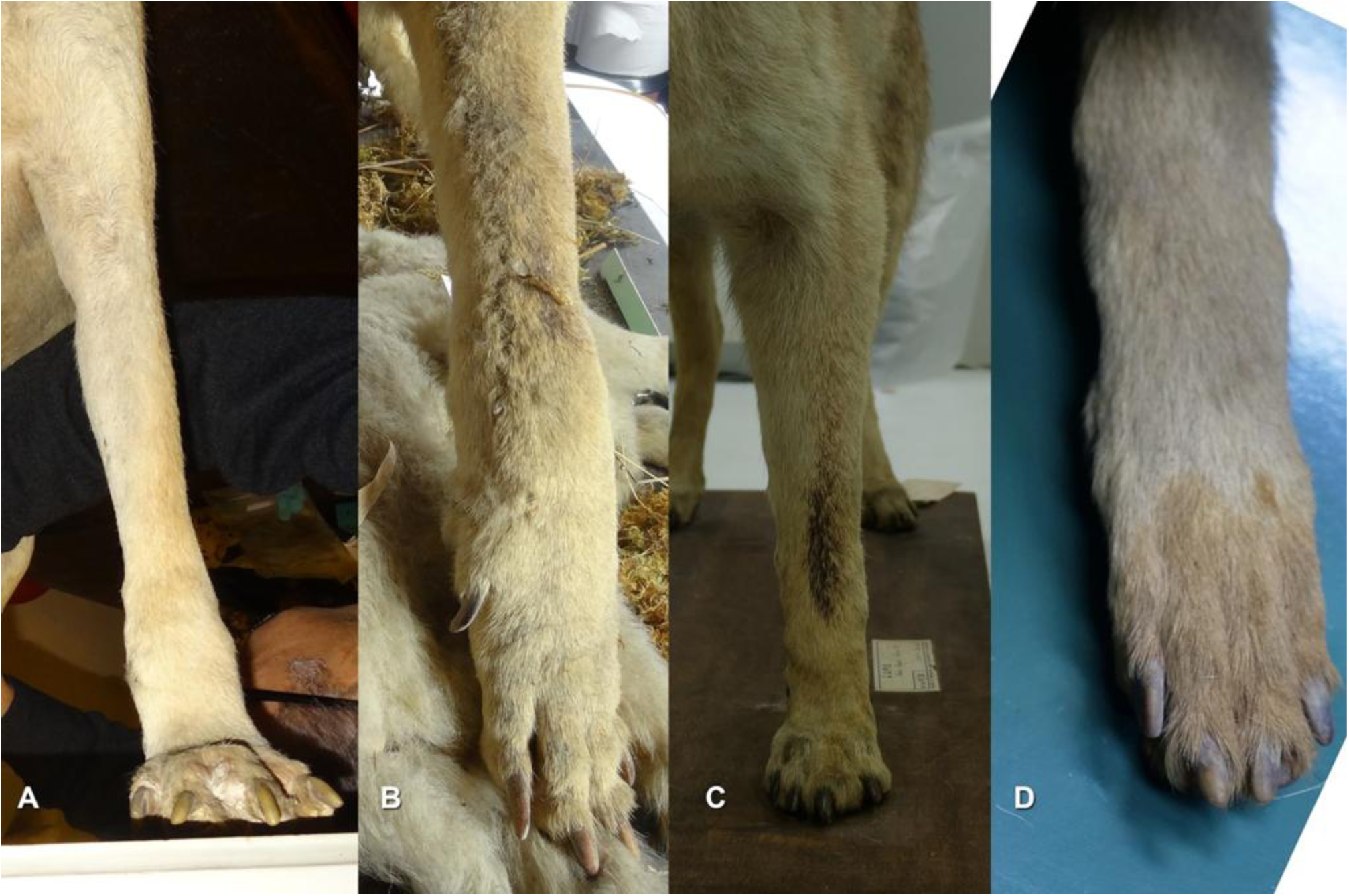
Forearms of Sicilian wolf have not, or just slightly hinted, line typical of Apennine wolf. A holotype, B paratype N.1, C paratype N. 2, D paratype N. 3.

The skull of the adult is smaller than the nominate subspecies and also significantly smaller than the Apennine subspecies *C. l. italicus*.

Comparison with Apennine wolf (*Canis lupus italicus* Altobello, 1921)

Table III shows the mean measurements of *Canis lupus cristaldii* subsp. nov., compared with those of *Canis lupus italicus* by Altobello (1921), and Ciucci and Boitani (2003)

**Table III.** Comparison between body measurements (mean) of *Canis lupus cristaldii* subsp. nov., and *Canis lupus italicus*.

|  | **Mean measures taken on *Canis lupus cristaldii* subsp. nov. (n= 4)** | **Mean measures taken on *Canis lupus italicus* (Altobello, 1921) (n >1)** | **Mean measures taken on *Canis lupus italicus* (processed from Ciucci and Boitani, 2003)** |
| --- | --- | --- | --- |
| Head-body length (cm) | 105.4 ($\pm 4.3$) | 127 | 109.1 ($\pm 6.18$, MM, n=18)<br>105.8 ($\pm 5.12$, FF, n=21) |
| Tail length (cm) | 34.9 ($\pm 5.9$) | 31 | 34.3 ($\pm 3.58$, n=27) |
| Shoulder height (cm) | 54.6 ($\pm 2.2$) | 65 | 66.9 ($\pm 7.66$, MM, n=12)<br>65.0 ($\pm 5.33$, FF, n=16) |
| Ear length (cm) | 8.4 ($\pm 1.1$) | 11 | 11.0 ($\pm 0.62$, n=24) |

The size of the Sicilian wolf is smaller than the average size of the Apennine wolf, in particular shoulder height, ear length, and head-to-body length.

The skull is also appreciably smaller than the skull of the Apennine wolf (Table IV). In the comparisons of Figs. 8–9 we used the skull of an adult female of Apennine wolf from Fiumicino di Premilcuore (FC), Appennino Tosco-Romagnolo, Central Italy (Museum of Ecology of Meldola, FC, labeled 194/2010).

**Table IV.**
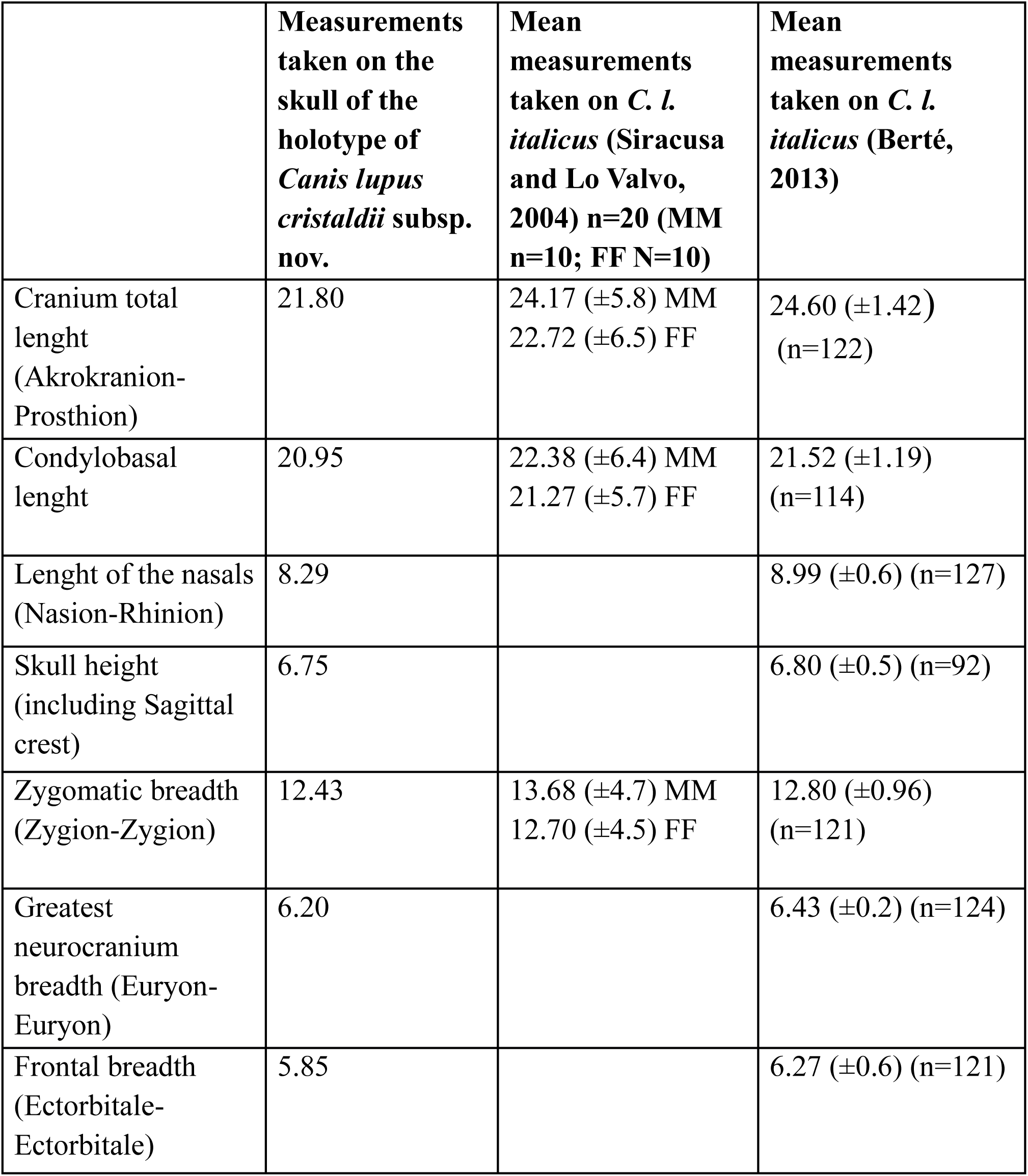
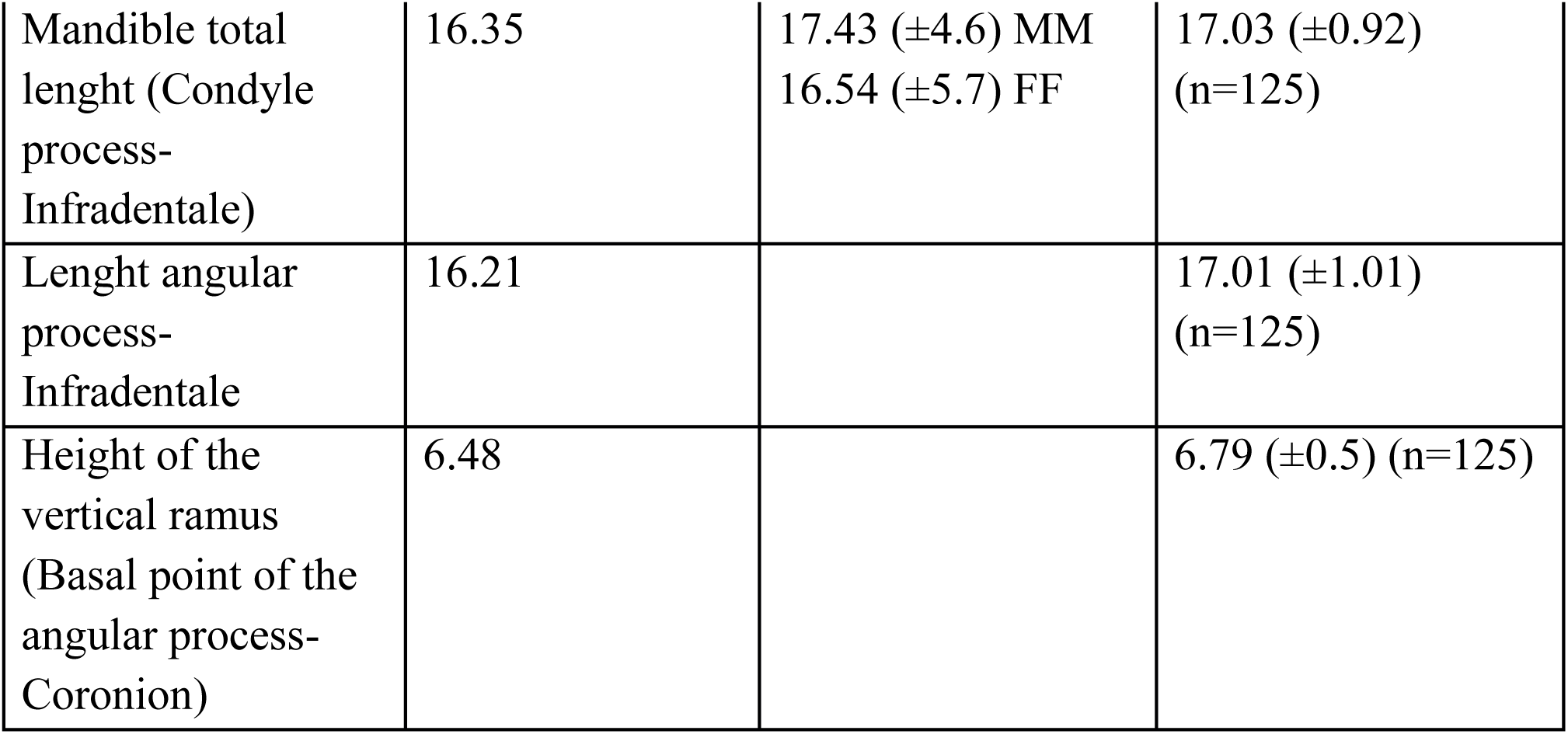
Measurements (cm) of the skull of the holotype of *Canis lupus cristaldii* **subsp. nov**. compared with *C. l. italicus* (Siracusa and Lo Valvo 2004; Berté 2013)

|  | **Measurements**<br>**taken on the**<br>**skull of the**<br>**holotype of**<br>***Canis lupus***<br>***cristaldii* subsp.**<br>**nov.** | **Mean**<br>**measurements**<br>**taken on *C. l.***<br>***italicus* (Siracusa**<br>**and Lo Valvo,**<br>**2004) n=20 (MM**<br>**n=10; FF N=10)** | **Mean**<br>**measurements**<br>**taken on *C. l.***<br>***italicus* (Berté,**<br>**2013)** |
| --- | --- | --- | --- |
| Cranium total<br>length<br>(Akrokranon-Prosthion) | 21.80 | 24.17 ($\pm 5.8$) MM<br>22.72 ($\pm 6.5$) FF | 24.60 ($\pm 1.42$)<br>(n=122) |
| Condylbasal<br>length | 20.95 | 22.38 ($\pm 6.4$) MM<br>21.27 ($\pm 5.7$) FF | 21.52 ($\pm 1.19$)<br>(n=114) |
| Length of the nasals<br>(Nasion-Rhinion) | 8.29 | | 8.99 ($\pm 0.6$) (n=127) |
| Skull height<br>(including Sagittal<br>crest) | 6.75 | | 6.80 ($\pm 0.5$) (n=92) |
| Zygomatic breadth<br>(Zygion-Zygion) | 12.43 | 13.68 ($\pm 4.7$) MM<br>12.70 ($\pm 4.5$) FF | 12.80 ($\pm 0.96$)<br>(n=121) |
| Greatest<br>neurocranium<br>breadth (Euryon-Euryon) | 6.20 | | 6.43 ($\pm 0.2$) (n=124) |
| Frontal breadth<br>(Ectorbitale-Ectorbitale) | 5.85 | | 6.27 ($\pm 0.6$) (n=121) |
| Mandible total length (Condyle process-Infradentale) | 16.35 | 17.43 ($\pm 4.6$) MM<br>16.54 ($\pm 5.7$) FF | 17.03 ($\pm 0.92$) (n=125) |
| Length angular process-Infradentale | 16.21 | | 17.01 ($\pm 1.01$) (n=125) |
| Height of the vertical ramus (Basal point of the angular process-Coronion) | 6.48 | | 6.79 ($\pm 0.5$) (n=125) |

**Fig. 8.**
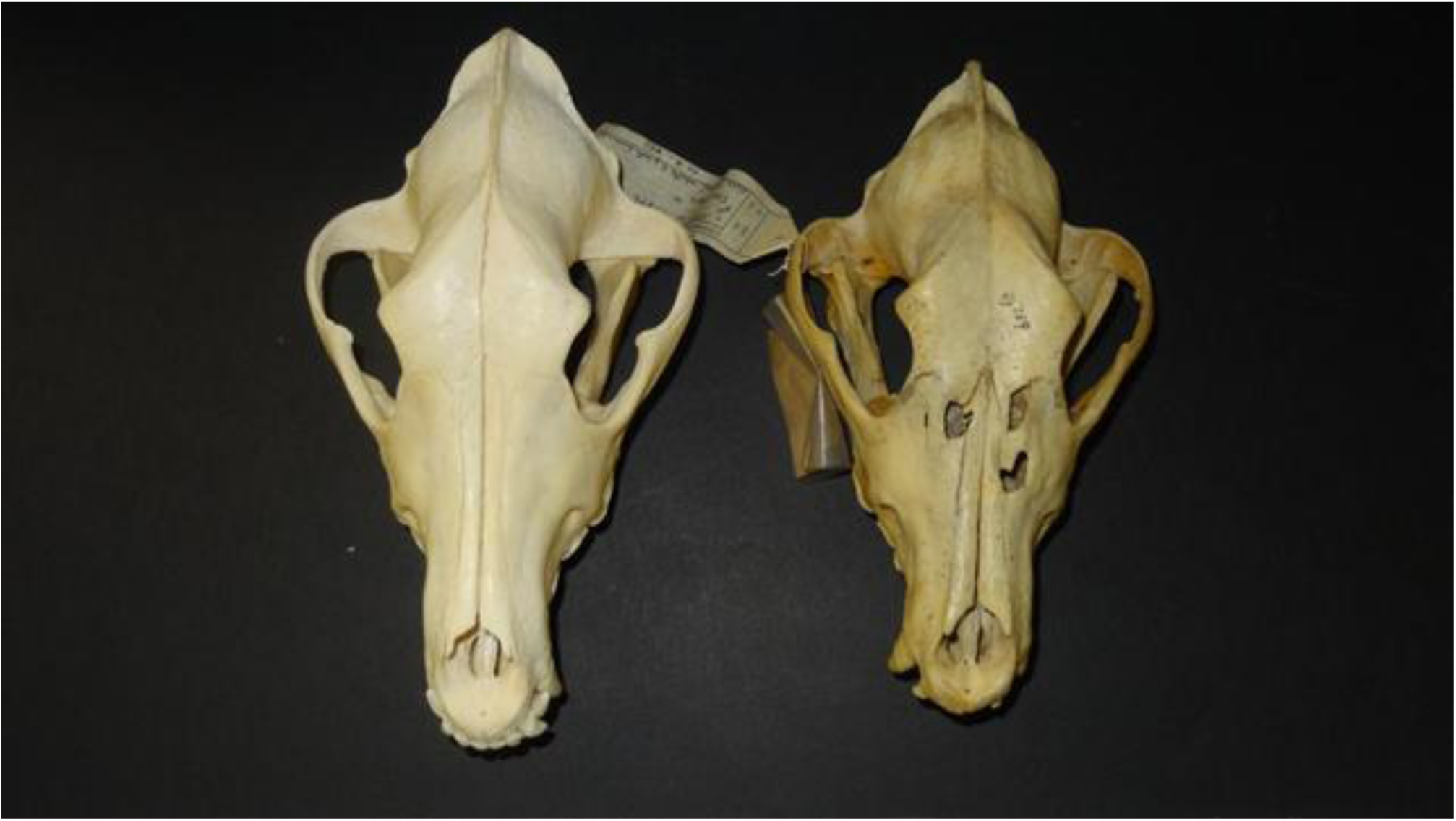
Size comparison between skulls of *Canis lupus cristaldii* subsp. nov. Holotype (right), and *Canis lupus italicus* (female) (left). For details see text.

**Fig. 9.**
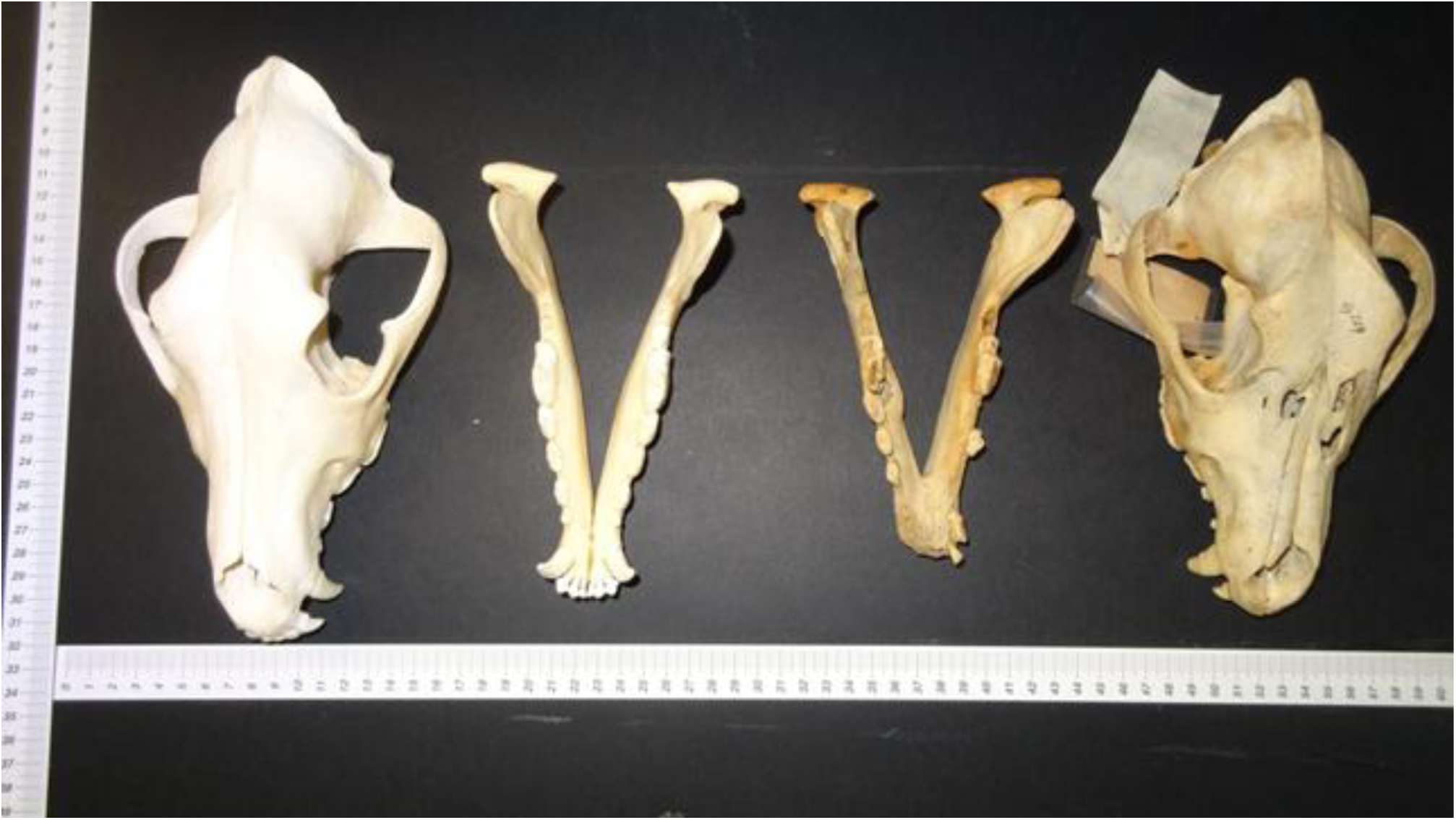
Size comparison between skulls and mandibles of *Canis lupus cristaldii* subsp. nov. Holotype (right), and *Canis lupus italicus* (female) (left). For details see text.

Fig. 10 shows the PCA of the body measurements of *C. l. italicus* (Ciucci & Boitani 2003) compared with four individuals of C. l. cristaldii susp. nov. In Fig. 11 is the correlation circle as explanation of Fig. 10 where is indicated how much they vary between the two comparative samples and how much they affect, the four linear measures collected. While in Fig. 12 is the dendrogram showing the phenetic relationship of the same samples of Fig. 10.

**Fig. 10.**
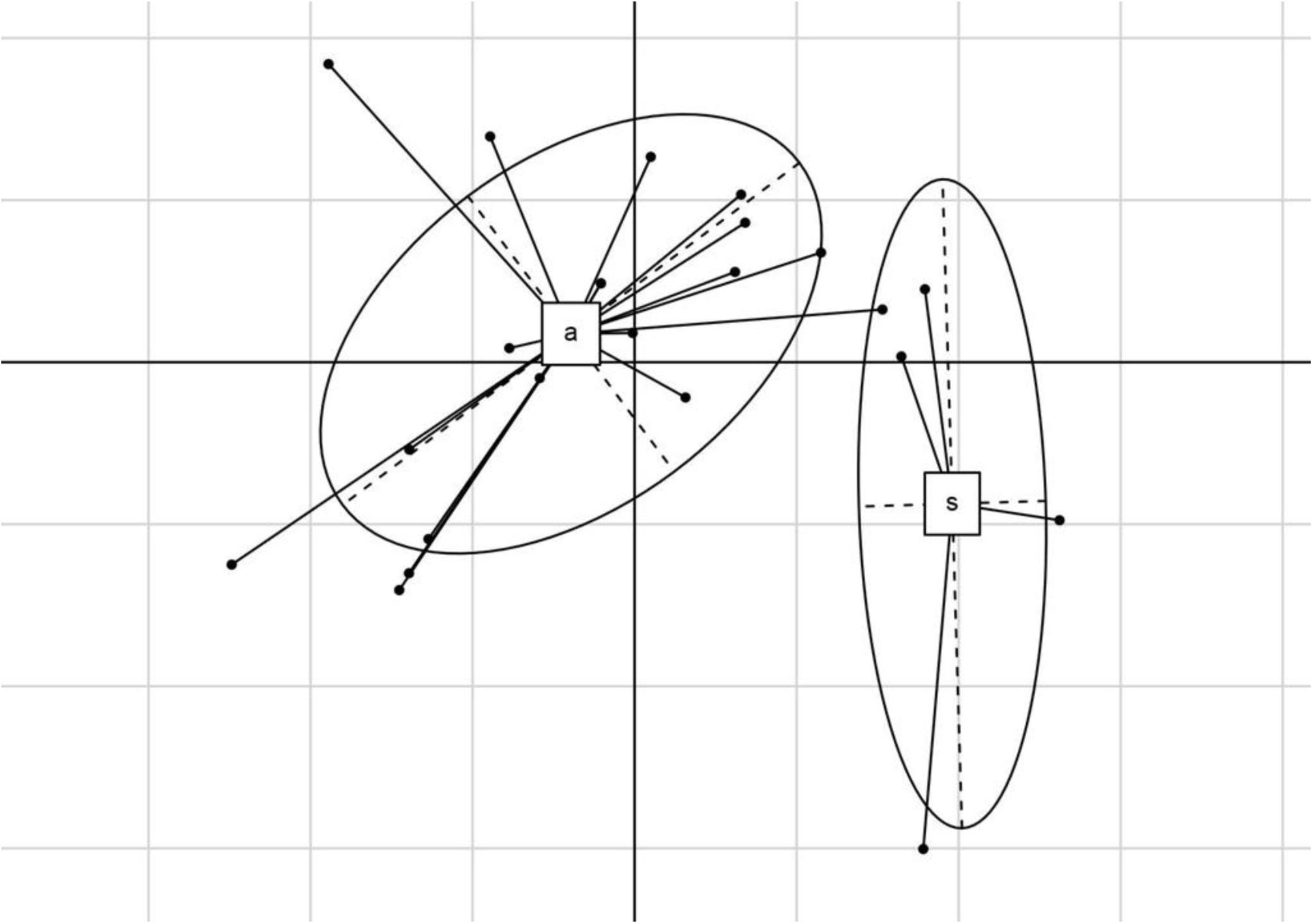
Principal Component Analysis (PCA) of the body measurements of *Canis lupus italicus* (a), compared with *Canis lupus cristaldii* subsp. nov. (s).

**Fig. 11.**
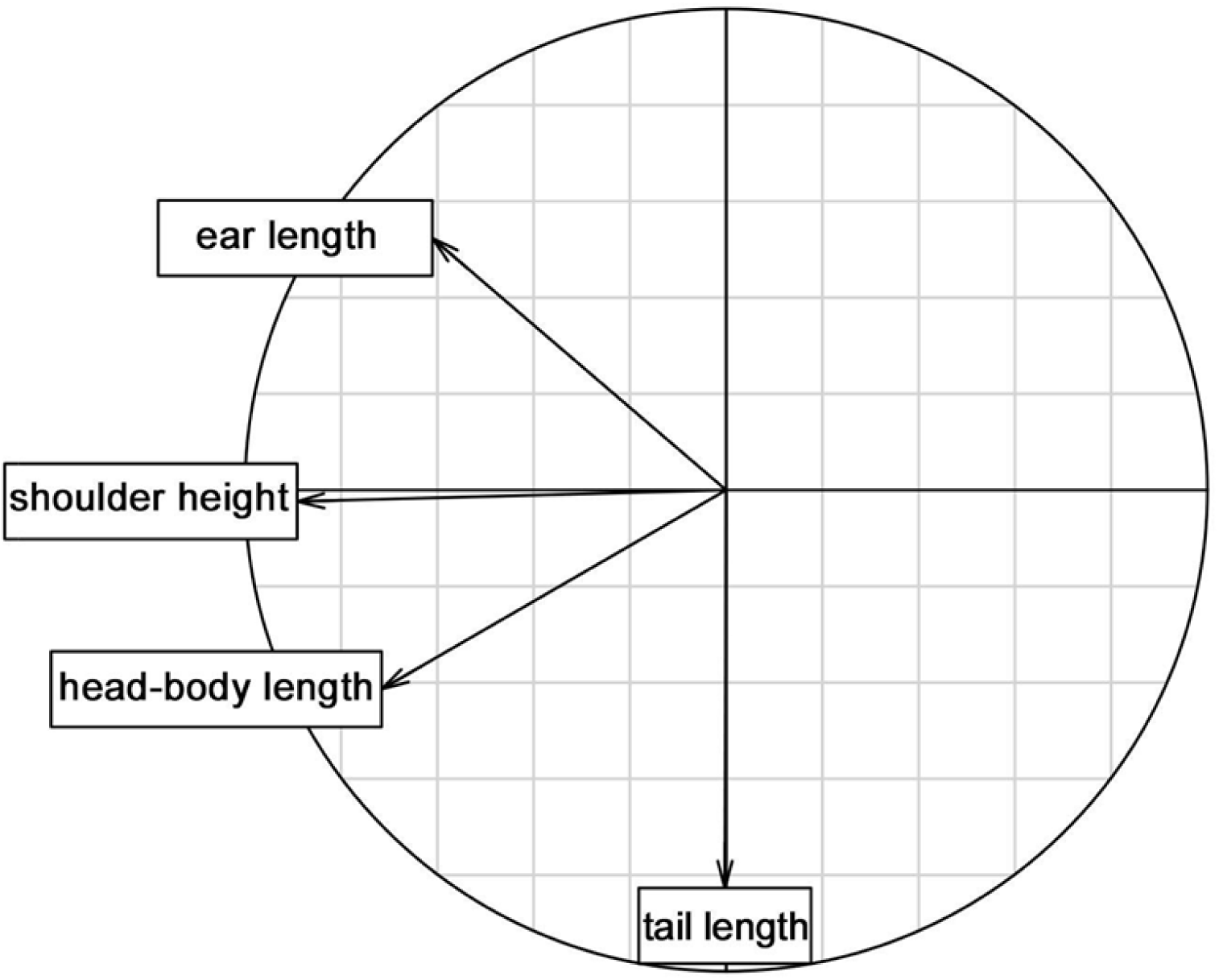
Correlation circle as explanation of Fig. 10. In this figure is indicated as the four linear measurements of the wolf body vary along the Cartesian axes, and how much they influence the separation of the two wolf samples

**Fig. 12.**
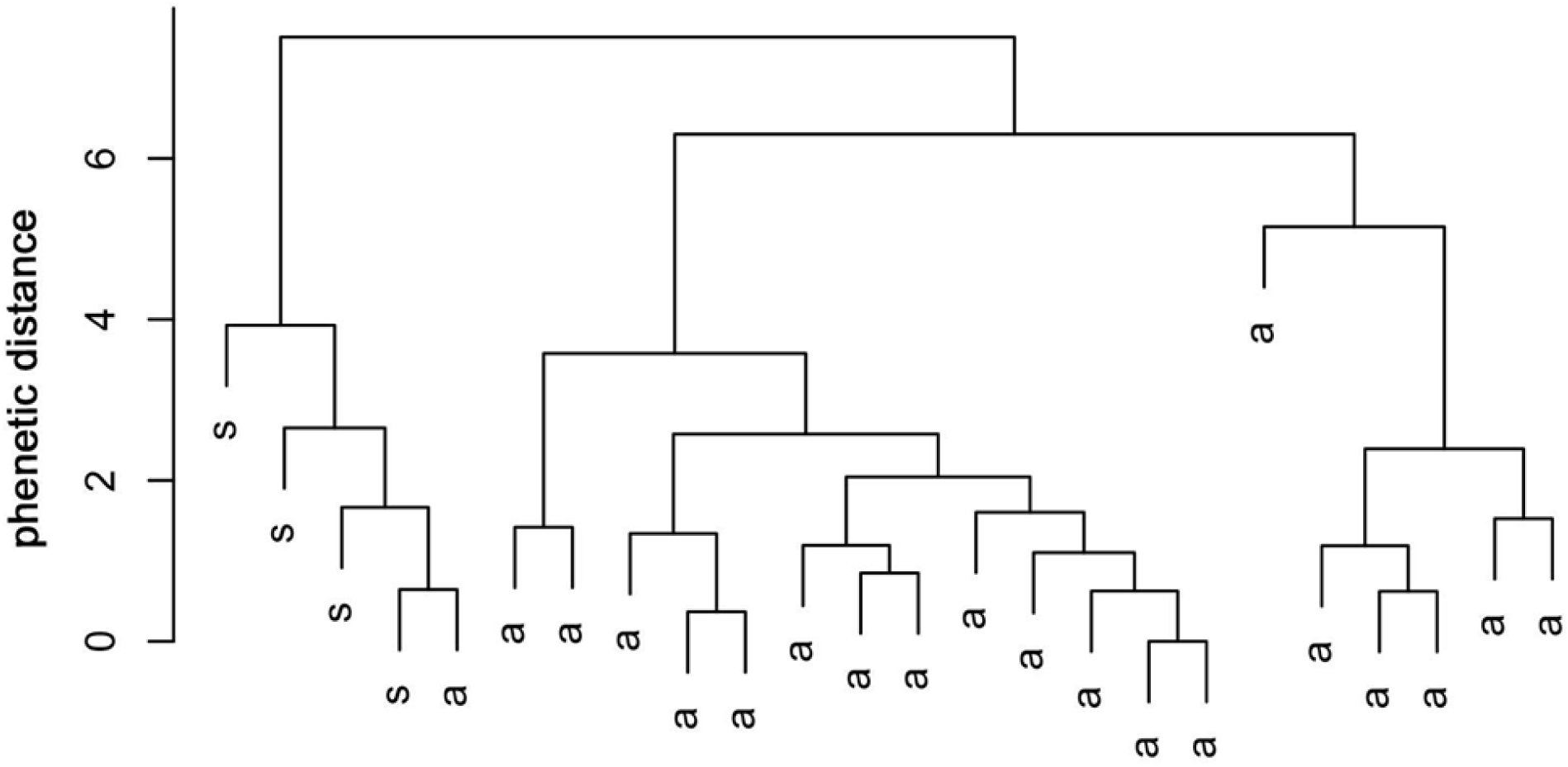
Dendrogram showing the phenetic relationships of the same samples of Fig. 10.

## Discussion

Data show that both holotype, and paratypes of *C. l. cristaldii* subsp. nov. have distinct phenotypic features from the Apennine wolf.

Fig. 10 shows the body measurements of four specimens of *C.l. cristaldii* subsp. nov. compared with the same body measurements of the sample of *C.l. italicus* in a two-dimensional space. The figure shows body measurements of the Sicilian wolf are clearly different from the Apennine wolf. Shoulder height is the trait that most differentiates Sicilian wolf from Apennine wolf (see Fig. 11). Ear length is also a notable trait, and also the head-body length, albeit less so. Tail length, however, appears to be exactly the same for both subspecies. Fig. 12 shows the Sicilian wolf to be readily separated from a phenetics point of view (ordinates) from the Apennine wolf because of its small size. In fact only one individual of *C.l. italicus* appears stochastically partially associated with *C.l.cristaldii*.

The morphological distinction is supported by preliminary analysis of mitochondrial DNA (mtDNA) which was conducted using genetic material extracted from the teeth of two specimens: the holotype skull, and an immature wolf skull preserved at the Museo di Zoologia ‘P. Doderlein’, Università di Palermo. The result indicates that both specimens to not belong to the Apennines population from mainland Italy because they present a unique and clearly differentiated haplotype which differs from the typical haplotype of the Apennine wolf (W14) by two substitutions (Angelici *et al*. 2016b). Further and more detailed analysis of mtDNA and genome sequencing are currently underway.

At the present state of knowledge we believe that the most likely explanation for these differences is the isolation from the Italian population from which the Sicilian wolf originated. The last known land bridge between Italy and Sicily is estimated between 21.5 and 20 kiloyears BP (Antonioli *et al*. 2012). Although the Sicilian mammal fauna is still not as well studied taxonomically, small mammals are increasingly recognized as discreetly endemic.(cf. Bezerra *et al*. 2016).

## Acknowledgments

This work would not have been possible without the heartfelt collaboration and generous help of many people. Paolo Agnelli of the Museo di Storia Naturale ‘La Specola’ helped us obtain continuous access to the holotype.

Ferdinando Maurici and Fabio Lo Valvo of the Museo Regionale Interdisciplinare ‘D’Aumale’ of Terrasini were just as generous and indispensable, as well as Sabrina Lo Brutto, Enrico Bellia, Maurizio Sarà of the Museo Zoologico ‘P. Doderlein’ of Palermo, and Fabio Lo Bono of the Museo Civico ‘B. Romano’ of Termini Imerese. A huge thank you goes to all of them.

We thank Agatino Maurizio Siracusa, Elisabetta Cilli, Marta M. Ciucani, Davide Palumbo, Romolo Caniglia, Riccardo Castiglia, Elena Fabbri, Flavia Annesi from the Sicilian wolf biomolecular working group for the advice and collaboration; E. Cilli was indispensable in pointing out the existence of the stuffed specimen of the museum of Termini Imerese. A. M. Siracusa has also collected some valuable historical bibliographic data. Davide F. Berté have made available to us many of his unpublished data on *C. l. italicus*, from his Ph.D thesis. A particular thank to Paolo Colangelo for the indispensable help in the statistical analysis, and his advice in compiling the present work. Our thanks also go to Paolo Laghi and Giancarlo Tedaldi of Museo di Ecologia di Meldola, for giving us the skull of female Apennine wolf for comparative pictures. Luigi Boitani and Paolo Ciucci have provided us with the database of the body size of the Apennine wolf. We are very grateful to Ronald H. Pine because his suggestions have been fundamental to increase the quality of the manuscript. We express our gratitude to Mauro Cella for improving the English of this manuscript.

## References

Altobello, G. (1921) Mammiferi. IV. I Carnivori (Carnivora). Fauna dell’Abruzzo e del Molise. Colitti, Campobasso, 61 pp.

Angelici, F.M., Rossi, L. & Siracusa, A.M. (2016a) The grey wolf in Sicily: a short history of an extinction. *In*: Angelici, F.M. & Rossi, L. (Eds). Atti del III Congresso Nazionale Fauna Problematica (Cesena, 24-26 November 2016). Cesena, pp. 99–100.

Angelici, F.M., Angelini, S., Annesi, F., Castiglia, R., Cilli, E., Ciucani, M.M., Ravegnini, G., Rossi, L. & Siracusa A.M. (2016b) Genetic identity of Sicilian grey wolf (*Canis lupus*) through Ancient DNA analyses. *In*: Angelici, F.M. & Rossi, L. (Eds). Atti del III Congresso Nazionale Fauna Problematica (Cesena, 24-26 November 2016). Cesena, pp. 108–109

Antonioli, F., Lo Presti, V., Morticelli, M.G., Mannino, M.A., Lambeck, K., Ferranti, L., Bonfiglioli, C., Mangano, V., Sannino, G.M., Furlani, S. & Canese, S.P. (2012) The land bridge between Europe and Sicily over the past 40 kyrs: timing of emersion and implications for the migration of *Homo sapiens*. Rendiconti Online della Società Geologica Italiana, 21: 1167–1169.

Berté, D.F. (2013) L’evoluzione del genere Canis (Carnivora, Canidae, Caninae) in Italia dal wolf-event a oggi: implicazioni biocronologiche, paleoecologiche e paleoambientali. Tesi di dottorato, XXVI ciclo. Sapienza Università di Roma, Dipartimento di Scienze della Terra, 390 pp.

Bezerra, A.M.R., Annesi, F., Aloise, G., Amori, G., Giustini L. & Castiglia R. (2016) Integrative taxonomy of the Italian pine voles, *Microtus savii* group (Cricetidae, Arvicolinae). Zoologica Scripta, 45: 225–236.

Bresc, H. (1983) “Disfari et perdiri li fructi et li aglandi”: economie e risorse boschive nella Sicilia medievale (XIII-XV secolo). Quaderni Storici, 18, 54 (3): 941–969.

Ciucci, P. & Boitani, L. (2003) Canis lupus, Linnaeus, 1758. *In*: Boitani, L., Lovari, S. & Vigna Taglianti, A. (Eds.). Fauna d’Italia. Mammalia III. Carnivora—Artiodactyla. Calderini, Bologna, pp. 20–47.

Chicoli, N. (1870). L’allevatore degli animali domestici in Sicilia. Seconda parte. Zootecnia speciale. Memorie scientifiche premiate per il concorso dal Congresso Agrario di Agrigento nel 1869. Stamp. Giovanni Lorsnaider, Palermo.

Galvagni, G.A. (1837). Fauna Etnea o materiali per la compilazione della zoologia dell’Etna. Memoria terza sulla terza famiglia dei dilaniatori. I Carnivori. Atti dell’Accademia Gioenia di Scienze Naturali di Catania, 13: 163–205.

La Mantia, T. & Cannella, Z. (2008) Presenza storica dei grossi Mammiferi in Sicilia. *In*: VV.AA., (2008) Atlante della Biodiversità della Sicilia: Vertebrati terrestri. Studi e Ricerche, 6. Arpa Sicilia, Palermo, pp. 87–112.

Mech, L.D. & Boitani, L. (2010) Canis lupus. The IUCN Red List of Threatened Species 2010: e.T3746A10049204. Available from: http://dx.doi.org/10.2305/IUCN.UK.2010-4.RLTS.T3746A10049204.en (December 2017)

Migneco, M. (1897) Considerazioni ed appunti sul cane cirneco. Stabilimento tipografico M. Galati, 17 pp.

Minà Palumbo, F. (1858a). Storia naturale delle Madonie. Catalogo dei Mammiferi. La Scienza e la Letteratura, A. I, 3: 154–170.

Minà Palumbo, F. (1858b) Storia naturale delle Madonie. Osservazioni sopra i Mammiferi. La Scienza e la Letteratura, A. I, 4: 5–14.

Minà Palumbo, F. (1868) Catalogo dei mammiferi della Sicilia. Annali di Agricoltura Siciliana, 12 (serie 2): 3–123.

Montana, L., Caniglia, R., Galaverni, M., Fabbri, E. & Randi, E. (2017) A new mitochondrial haplotype confirms the distinctiveness of the Italian wolf *(Canis lupus)* population. Mammalian Biology, 84: 30–34.

Nowak, R.M. & Federoff, N.E. (2002) The systematic status of the Italian wolf *Canis lupus*. Acta Theriologica, 47: 333–338. doi:10.1007/bf03194151

Pocock, R.I. (1935) The races of *Canis lupus*. Journal of Zoology, 105: 647–686.

Siracusa, A.M. & Lo Valvo, M. (2004) Confronti craniometrici tra lupi (*Canis lupus*) dell’Italia continentale e della Spagna: primi dati. Hystrix, Italian Journal of Mammology, 15: 31–38.

Von den Driesch, A. (1976) A guide to the measurement of animal bones from archaeological sites. Peabody Museum Bulletin, 1. Harvard. pp. 136.

Ward, J. H. Jr., (1963) Hierarchical Grouping to Optimize an Objective Function. Journal of the American Statistical Association, 58: 236–244.

Wozencraft, W.C, (2005) Order Carnivora. *In*: Wilson, D.E. & Reeder, D.A.M. (Eds), Mammal Species of the World: A Taxonomic and Geographic Reference (3rd ed.). Johns Hopkins University Press, Baltimore, Maryland pp. 532–628.

